# Testing frameworks for early life effects: the developmental constraints and adaptive response hypotheses do not explain key fertility outcomes in wild female baboons

**DOI:** 10.1101/2024.04.23.590627

**Authors:** Stacy Rosenbaum, Anup Malani, Amanda J. Lea, Jenny Tung, Susan C. Alberts, Elizabeth A. Archie

## Abstract

In evolutionary ecology, two classes of explanations are frequently invoked to explain “early life effects” on adult outcomes. Developmental constraints (DC) explanations contend that costs of early adversity arise from limitations adversity places on optimal development. Adaptive response (AR) hypotheses propose that later life outcomes will be worse when early and adult environments are poorly “matched.” Here, we use recently proposed mathematical definitions for these hypotheses and a quadratic-regression based approach to test the long-term consequences of variation in developmental environments on fertility in wild baboons. We evaluate whether low rainfall and/or dominance rank during development predict three female fertility measures in adulthood, and whether any observed relationships are consistent with DC and/or AR. Neither rainfall during development nor the difference between rainfall in development and adulthood predicted any fertility measures. Females who were low-ranking during development had an elevated risk of losing infants later in life, and greater change in rank between development and adulthood predicted greater risk of infant loss. However, both effects were statistically marginal and consistent with alternative explanations, including adult environmental quality effects. Consequently, our data do not provide compelling support for either of these common explanations for the evolution of early life effects.

## 1 Introduction

In many animal species, including humans, exposure to early life socioecological adversity predicts a wide range of negative outcomes in adulthood, including poor health, reduced fitness, compromised social functioning, and shorter lifespans [1, 2, 3, 4, 5, 6, 7]. Given the diverse range of species in which such “early life effects” are observed, there is considerable interest in understanding the evolutionary origins of the connections between early life experiences and later life outcomes [8, 9, 10, 11, 12, 13, 14].

Two major classes of conceptual models are commonly invoked to explain early life effects in the evolutionary ecology literature [15, 16], and have influenced thinking about the evolution of early life effects in humans [9, 12, 17]. The first class, developmental constraints (DC, also known as silver spoon) hypotheses, proposes that early life adversity leads to morphological, physiological, and/or behavioral tradeoffs that prioritize short-term survival but carry long-term costs, such as shortened lifespans or poor health in adulthood [8, 15]. The second class, which we will refer to collectively here as adaptive response (AR) hypotheses, proposes that organisms adopt a phenotype suited to either their developmental environment (the developmental adaptive response hypothesis) or to their predicted future environment (the predictive adaptive response hypothesis, including both “internal” and “external” predictive adaptive responses) [18, 19, 20]. In either scenario, they incur fitness costs when the phenotype they adopt does not suit the environment they find themselves in during adulthood. Though they are sometimes framed as alternative explanations, the two classes of models do not have to be mutually exclusive [21, 22, 23, 24]. Developmental constraints hypotheses generate predictions about the downstream effects of an organism’s “starting point” (i.e., the quality of their early life environment), while adaptive response hypotheses generate predictions about the relationship between environmental stability across the lifespan and adult outcomes.

In animals with slow life histories, there is more empirical support for developmental constraints than for adaptive response hypotheses [reviewed in 14, 16, 25]. Many studies have found that organisms do worse on a variety of fitness-relevant outcomes when they experience poor-quality developmental environments [6, 26], something we examine in more detail in the discussion (Section 4). However, it is difficult to know which hypothesis (or hypotheses) best explain this relationship. The intertwined nature of the variables under consideration complicates empirical tests: early life conditions and the difference between early life and adult conditions are not independent of one another. Consequently, commonly applied tests to differentiate the hypotheses may be vulnerable to high error rates and conflate DC and AR [27].

To help remedy this problem, we recently published formal (mathematical) definitions of developmental constraints, developmental adaptive response, and predictive adaptive response hypotheses and proposed empirical tests of these hypotheses derived from the definitions (Table 1, following [27]). Our definition of the developmental constraints model states that experiencing a worse developmental environment (*e*_0_) leads to worse outcomes in adulthood (*y*). Our definition of the developmental adaptive response model states that organisms alter their phenotype to adapt to their developmental environment (i.e., to *e*_0_), while our definition of the predictive adaptive response model states that organisms adapt their phenotype in anticipation of a predicted future adult environment (*e*_1_). Both AR definitions specifically posit that adult outcomes are a function of the difference between developmental and adult environments, and arise as a consequence of the organism’s “choice” of a phenotype (*p*) based on the cues they receive from the early environment. Consequently, a primary prediction of the two AR hypotheses is that larger differences between the developmental and adult environments–i.e., larger environmental deltas (Δ*e*)–lead to worse outcomes in adulthood (Table 1). All three definitions are agnostic as to how, mechanistically, the connection between developmental environment and adult outcomes occurs.

**Table 1:**
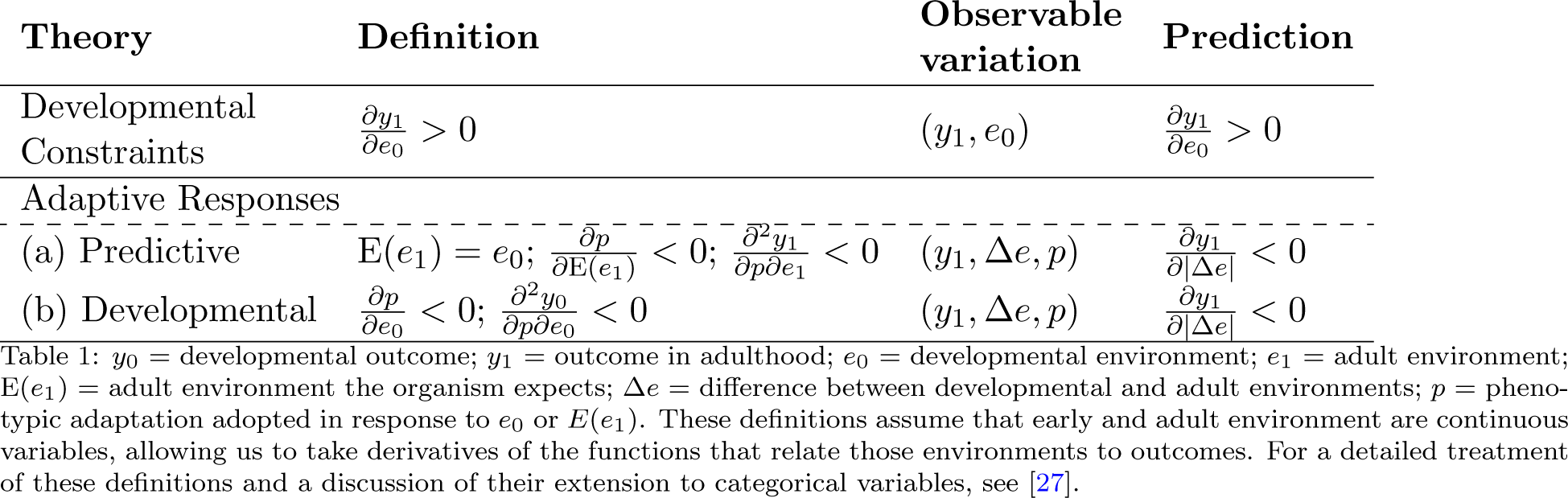
Formal definitions and derived prediction of theories for the relationship between the quality of developmental environments and adult outcomes.

In practice, the developmental and predictive adaptive response hypotheses can be difficult to distinguish empirically [16, 28, 29]. One common strategy for operationalizing the underlying phenomenon that constitutes an organism’s “prediction” is to assume that organisms predict that some feature(s) of their developmental environment (e.g., the amount of rainfall they experience or the size of the social group they live in) will be similar to what they experience as adults [12, 17, 30, 31]. If this prediction is correct (i.e., organisms predict accurately, and thus experience little difference between their developmental and adult environments, corresponding to a low Δ*e* in Table 1), then their outcomes will be better than if their prediction was incorrect (i.e., experience large differences between their developmental and adult environments, a high Δ*e*). However, the developmental adaptive response hypothesis makes the same basic prediction. Namely, it predicts that organisms will fare worse if they find themselves in an adult environment that is not suited to the phenotype they adopted in response to their developmental environment. Because the two hypotheses cannot be distinguished using measures of the environmental delta and adult outcomes alone [16, 27, 29], here we group them together as a single “adaptive response” (AR) hypothesis.

While we originally proposed our formal definitions and a strategy for empirical analysis in [27], we did not previously apply them to real data. Here, we do so for the first time by evaluating the evidence for DC and AR in wild female baboons monitored by the Amboseli Baboon Research Project (ABRP) [32]. The Amboseli baboons are excellent subjects for this research because they live in a highly variable environment that can generate considerable differences in the ecological and social environments experienced by the same animal across the course of their lives [4, 32, 33]. Additionally, because this population has been prospectively studied for multiple generations, now spanning more than 50 years, the types of data necessary to investigate the long-term effects of early life are available. Notably, relationships between early life experiences and later life outcomes, including life history traits like fertility and mortality, are also well-established for this study system [e.g., 4, 34, 35, 36, 37]. In social animals, aspects of both early life ecological and social environments may play important roles in determining later life outcomes, though the two are often difficult to tease apart given that social partners also represent competition for resources provided by the environment [16, 38, 39]. Here we examine the effects of two key socioecological variables that play important roles in outcomes for female baboons [4, 34, 40, 41, 42]. The first variable is dominance rank. Female baboons have linear dominance hierarchies with a strong pattern of non-genetic matrilineal rank inheritance [43, 44]. This means that many female baboons hold a rank in adulthood that is similar to the rank their mother held when they themselves were born. However, due to group fissions and matriline overthrows (where the female members of one family successfully challenge the females of a higher-ranked family, leading to them changing positions in the dominance hierarchy), some animals end up considerably higher or lower-ranking than the one they experienced (via their mother) during development [32, 44]. The other variable we consider is rainfall. The Amboseli ecosystem is semi-arid and highly seasonal, with large (more than four-fold) variance in annual rainfall totals [45, 46]. This means that throughout her adulthood, a female might experience years in which rainfall was similar to what she experienced during her first year of life, and years in which it was very different.

We capitalize on these characteristics of baboon socioecology to test the DC and AR hypotheses. Specifically, we first test the DC hypothesis by asking whether experiencing low rainfall or being low-ranking during development predicts lower odds of conceiving, giving birth to a live infant, and/or raising an infant to weaning age as an adult. Next, we test the AR hypothesis by asking whether larger dominance rank or rainfall deltas between development and adulthood predict these same three outcomes. Our data enable us to investigate the relative evidence for AR and DC in one of the longest-running field studies of wild social mammals to date, drawing on a theory-aligned statistical method to disentangle the relationship between developmental environments and environmental deltas on key fertility measures.

## 2 Methods

### 2.1 Study subjects

Our subjects were 295 wild female baboons in the hybrid population that resides in the Amboseli ecosystem in southern Kenya. Baboons in this population have primarily yellow baboon (*Papio cynocephalus*) ancestry, but also near-universal minority ancestry from Anubis baboons [47, 48]. This population has been studied since 1971 by the Amboseli Baboon Research Project (ABRP) [32], which collects longitudinal demographic, ecological, life history, and behavioral data on individually recognized animals. Reproductive state (e.g., cycling, pregnant), the timing of events (e.g., births, conceptions), and females’ ages were known based on direct, nearly-daily observations of females done by experienced ABRP researchers. Data included in the present analyses span the years of 1974 to 2023.

### 2.2 Outcome variables

We used three measures of fertility as outcome variables in our models. The first outcome was whether or not a female **conceived in a given observation month**, conditional upon entering the month in a reproductive state in which conception was possible (i.e., she was cycling on the first day of the month, not pregnant or in postpartum amenorrhea). The second outcome was whether or not a female **gave birth to a live infant**, conditional upon being recorded as pregnant (note that we miss many pregnancies that end in early-term miscarriages [49]. The third outcome was whether a female **successfully raised an infant to 70 weeks of age** (the average age at weaning [50]), conditional upon having successfully given birth to a live infant (Tables 2 & 3). Sample sizes differ across outcome variables because there are inevitably more conceptions than live births and more live births than infants who live to 70 weeks. Sample sizes in the rainfall models (Table 2) are slightly smaller than sample sizes in the models that evaluate rank (Table 3; e.g., n=1,154 instead of 1,158 conceptions) because rainfall data were not collected until 1976 and our rank data set begins in 1974. Any pregnancy or infant survival data that were censored in the ABRP data set (e.g., infants who were still too young to have reached 70 weeks of age, or pregnancies that were still in progress) were dropped from the analyses. For the infant survival analysis, we also excluded any observations where the subject died before her infant, as early maternal loss strongly predicts subsequent infant death.

**Table 2:**
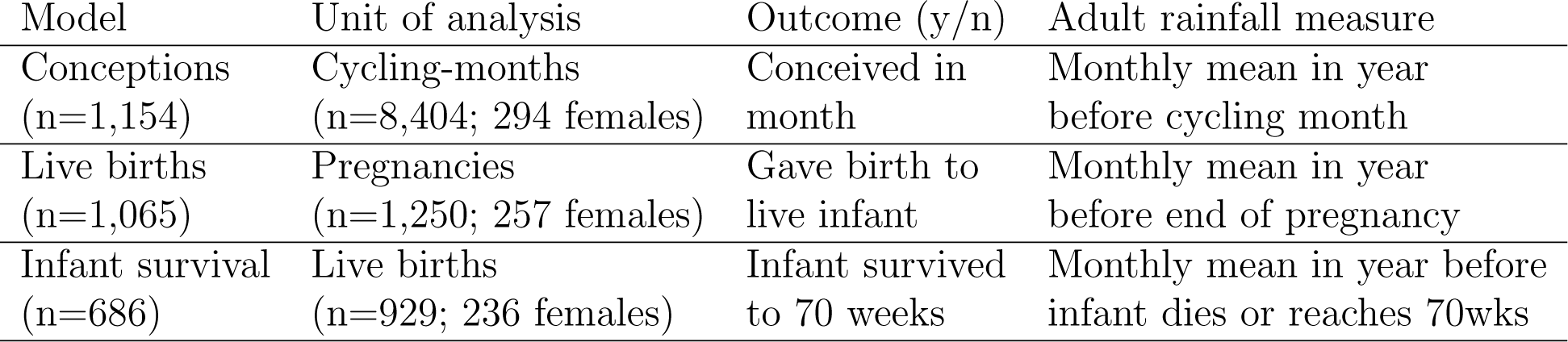
Basic descriptive information for conception, live birth, and infant survival data sets (rainfall analysis)

| Model | Unit of analysis | Outcome (y/n) | Adult rainfall measure |
| --- | --- | --- | --- |
| Conceptions<br>($n=1,154$) | Cycling-months<br>($n=8,404$; 294 females) | Conceived in<br>month | Monthly mean in year<br>before cycling month |
| Live births<br>($n=1,065$) | Pregnancies<br>($n=1,250$; 257 females) | Gave birth to<br>live infant | Monthly mean in year<br>before end of pregnancy |
| Infant survival<br>($n=686$) | Live births<br>($n=929$; 236 females) | Infant survived<br>to 70 weeks | Monthly mean in year before<br>infant dies or reaches 70wks |

**Table 3:**
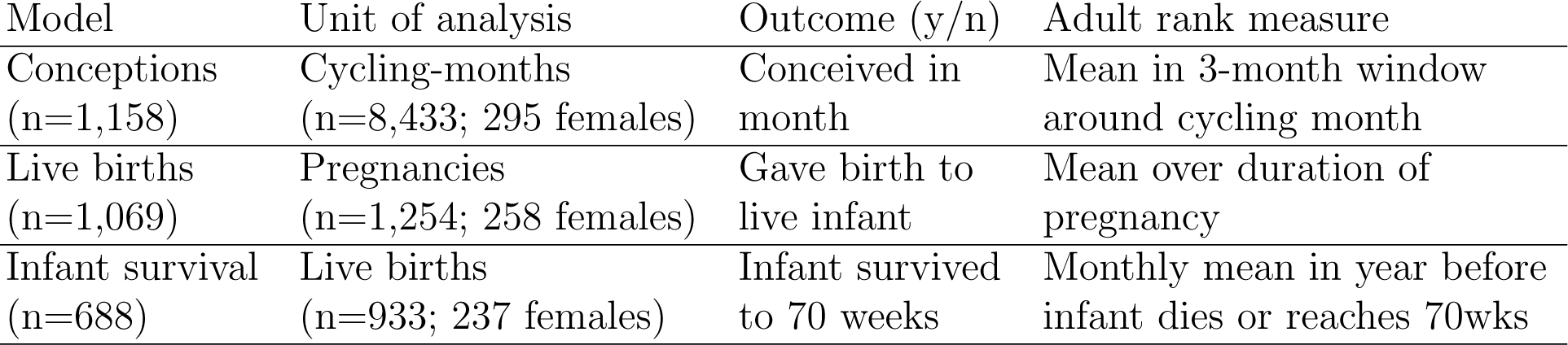
Basic descriptive information for conception, live birth, and infant survival data sets (dominance rank analysis)

### 2.3 Predictor variables

**Rainfall:** To test whether female baboons have better reproductive outcomes if they 1) are born in higher-rainfall years, and 2) experience smaller rainfall deltas (i.e., the rainfall they experienced in adulthood more closely “matches” the rainfall they experienced during development), we used precipitation data collected from a rain gauge that is checked daily at the ABRP field camp. For rainfall during development, we calculated average monthly rainfall in the subject’s first year of life. When averaged across the three data sets (Table 2), mean monthly rainfall during development was 28.80mm/month (SD=0.31, range=7.43-63.92). For rainfall in adulthood, we calculated average monthly rainfall in the 12 months before the fertility event (e.g. conception, birth) in question. Averaged across the three data sets (Table 2), mean monthly rainfall during adulthood was 30.53mm/month (SD=0.98, range=7.92-66.76). To calculate the rainfall deltas, we subtracted our measure of rainfall during development from our measure of rainfall in adulthood.

Histograms of the difference between rainfall during development and adulthood can be found in Figure 1 (top row) for all three data sets (conception: panel A; live birth: panel B; and infant survival: panel C, as described in Table 2). More detailed summary statistics concerning rainfall are available in Section A of the supplementary materials.

**Figure 1:**
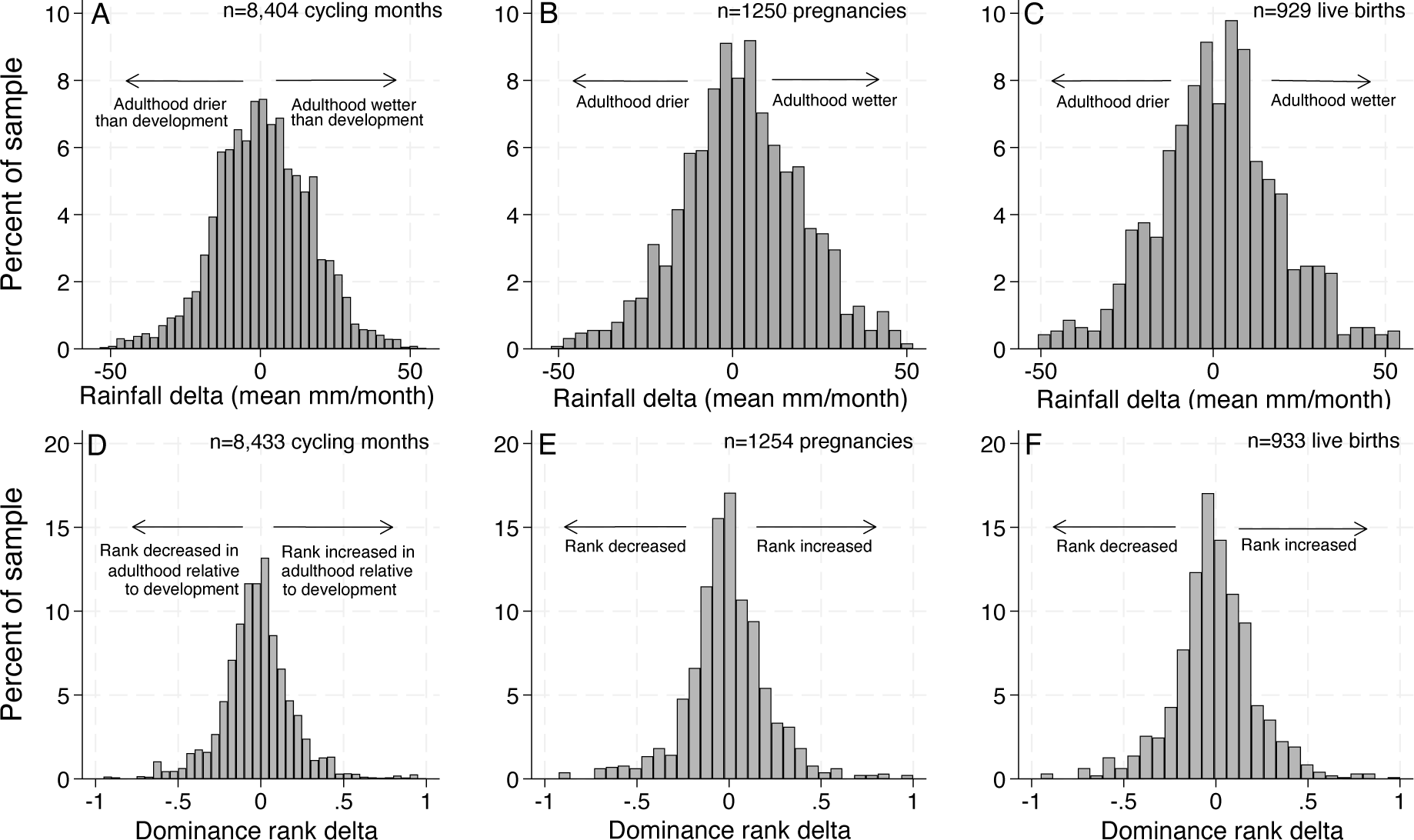
Difference between rainfall (top row) and dominance rank (bottom row) between development and adulthood. **Top row:** The distribution of rainfall deltas (i.e., the difference between the amount it rained during subjects’ first year of life and the amount it rained in the year of adulthood in which the outcome was measured (see Table 2)), expressed in mean mm/month. A value of zero on the x-axis indicates that it rained the same amount during development as it did during the year the fertility outcome was measured. Rainfall in adulthood was the average mm rain/month in the year before the fertility event in question (i.e., the 12 months before a given cycling month (Panel A); live birth or miscarriage (Panel B); or “termination” of an infant either because it died or because it successfully reached 70 weeks, the average age at weaning (Panel C). See Table 2 for a description of the three models corresponding to the three fertility outcome variables. **Bottom row:** The distribution of dominance rank deltas (i.e., the difference between subjects’ mothers’ proportional dominance rank when the subject was born, and the subjects’ proportional dominance rank in the period of adulthood in which the outcome was measured (see Table 3). The center of the x-axis (0) represents a perfect match between proportional rank at birth and adult rank when the outcome was measured. Negative x-axis values (bounded by −1) are subjects who held a lower rank in adulthood than their mothers did when they (the subjects) were born, and positive x-axis values (bounded by 1) are subjects who held a higher rank in adulthood than their mothers did when they (the subjects) were born. Rank during adulthood was the average rank of the subject in the three months centered around a given cycling month (Panel D), over the duration of the pregnancy (Panel E), or over the duration of the resulting infant’s life up to 70 weeks (Panel F), as appropriate for a given analysis. See Table 3 for a description of the three models corresponding to the three fertility outcome variables.

**Dominance rank:** To test whether female baboons have better reproductive outcomes if 1) they are higher-ranking during development, or 2) they have smaller rank deltas (i.e., their dominance rank in adulthood is similar to their rank during development) we used information about animals’ proportional dominance ranks. Proportional dominance rank represents the proportion of adult female group members that the subject in question outranks [51, 52, 53]. For example, a proportional dominance rank of 0.9 means that the female outranks 90% of the adult females in her group. Dominance was determined by the outcomes of all observed, decided agonistic interactions between individual females, and was estimated on a monthly basis [52, 53]. Additional details about how dominance rank data are collected and ranks assigned can be found in Section B of the supplementary materials and in [53].

Rank during development was defined as the average rank of the subject’s mother over the three-month span centering the subject’s birth (i.e., the month of the birth, plus the month before and the month after). Rank in adulthood was averaged over a three month window for the relevant potential cycling month (i.e., potential conception month); over the course of the pregnancy; or over the lifespan of the subject’s infant (up to 70 weeks) as appropriate for the respective outcome variable (Table 3). Using three month windows for rank during development and rank in a potential conception month allowed us to retain a slightly larger number of observations in the data set than using only a single month would, since rank data are occasionally missing for a female in a given month due to observation gaps. These gaps were rare: e.g., we missed the female subject’s rank during the focal month of observation for only 55 out of 8,433 cycling months in which rank data were available for her in either the month before, month after, or both. The correlation coefficient between average rank in the three month windows and rank strictly during the month of interest was *>* 0.99 for both rank during development and rank in adulthood. Histograms of the difference between rank during development and adulthood can be found in Figure 1 (bottom row) for all three data sets (conception: panel D; live birth: panel E; and infant survival: panel F, as described in Table 3).

#### 2.3.1 Covariates

Other variables besides those of interest here may influence female fertility, including age and group size (the latter because it may influence resource availability [34, 40, 54]). Thus, we included these known predictors of female fertility as covariates in our models. Age was operationalized as the female subject’s age at the start of the conception month, at pregnancy termination, and at infant birth, as appropriate for a given model (Tables 2 & 3). We also included a squared age term because the effects of age on fertility measures are non-linear: old and very young females tend to be less fertile [55]. Group size data were obtained via group censuses conducted during routine research visits. Group size was defined as the average number of group members each day during the cycling month, across the pregnancy, and across the resulting infant’s life, again as relevant for a given model. See Section C and Table A.4 in the supplementary materials for age and group size summary statistics, as well as more details about the measurement and operationalization of these covariates.

Prior Amboseli research has indicated that nulliparous females frequently require more cycling months to conceive than other females do (i.e., their odds are lower in any given month), and that females whose last infant died before weaning require fewer cycling months to conceive than females whose prior infant survived [54, 56, 57]. We chose not to include parity or the status of a prior infant as covariates in the conception models in the main text because we are unsure if rain or rank during development, or delta rain or rank, influence time until first conception or time until conception after an infant loss (i.e., contribute to the pathways through which these variables might have an effect). For example, if a female who experienced drought during development is more likely to lose her infant than females who did not experience drought, controlling for infant loss in models of conception could mask the relationship between drought during development and adult conception probability.

Nonetheless, for comparison purposes, in the supplement we present versions of the conception models that only include parous females whose prior infant lived to at least 70 weeks. That is, we exclude nulliparous females and females whose prior infant died before weaning. This decision reduces the sample size by 60%. There are qualitative differences in the results obtained from the larger data set and those obtained from this smaller subset of the data; specifically, in this reduced sample there is weak support for the mismatch prediction of AR. However, 1) the support only occurs in across-female and within-group versions of the models, which are less powerful tests of AR than the within-female version we use (discussed in detail in Section 2.4), and 2) the results do not approach statistical significance after multiple testing adjustments are applied. This result is fully presented in Section D and Table A.5 in the supplement.

### 2.4 Analysis strategy

Our formal versions of the DC and AR hypotheses posit that variation in the environment (either early life itself or the difference between early life and adult environments) is causally responsible for differences in later-life outcomes. The ideal test of the DC hypothesis would be to vary the early life environment of the animals while holding all else constant. Because this type of experiment is impossible in natural populations, we instead compare across individuals under the assumption that the environmental measures of interest (here, rank and rainfall) are independently distributed across our study subjects [58, 59].

In the case of rainfall this is likely a reasonable assumption, as no property of the baboons themselves influences how much it rains in Amboseli. In the case of rank, the assumption is more questionable (see discussion in Section 4): females of different ranks, or who fell, rose, or maintained the same rank are likely to be different from one another in other, unobserved ways. For example, a female whose rank fell might be more likely to be sick or in poor condition than a female whose rank stayed the same. A female whose rank rose significantly might be in especially good condition or may have been more likely to experience a group fission event in which she was able to move to the top of a new hierarchy. For the AR hypothesis, we can make both inter-and intra-individual comparisons because the same female can be observed multiple times in adulthood, during which her environment can change. For example, the same female baboon could experience drought in one year but plentiful rainfall in another, or change rank between years.

Our regression models and their associated tests are derived from the formal mathematical definitions of DC and AR provided in Table 1. Following [27], we use a quadratic regression model:

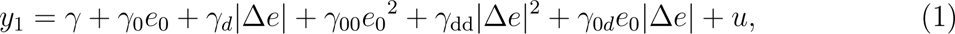

where *y*_1_ is an outcome in adulthood, *e*_0_ is the developmental environment, Δ*e* is the difference between the developmental and adult environments, the subscript 0 on *γ* indicates the coefficient is on *e*_0_, *d* indicates it is on *|*Δ*e|*, 0*d* indicates it is on *e*_0_*|*Δ*e|*, and 00 indicates that it is on *e*_0_^2^. The quadratic regression is useful because it is mathematically compatible with the idea that outcomes can get worse both when adult environments are worse than developmental environments *and* when they are better than developmental environments, which is a key prediction of many versions of the AR hypothesis.

A term for the quality of the adult environment is deliberately not included in our primary models. This is because of the non-independence of *e*_0_, Δ*e*, and adult environment (*e*_1_). If *e*_0_ is held constant and *e*_1_ is different than *e*_0_, then it is impossible to tell if any observed changes in the outcome are due to the value of *e*_1_ or to Δ*e*. Following [27], we assume that what best represents the biological phenomenon under consideration is that *e*_0_ and Δ*e* are independent variables that collectively generate *e*_1_ via the relationship *e*_0_ + Δ*e* = *e*_1_. We make this assumption because an organism does not experience *e*_1_ independent of what they experienced during *e*_0_. Because of this assumption [60], *e*_1_ is simply an endogenous byproduct of the structure of the theoretical model, and thus not a testable variable.

The above regression (equation 1) implies different statistical tests for DC and AR. Following the definition of DC given in Table 1, we test for DC using the following inequality:

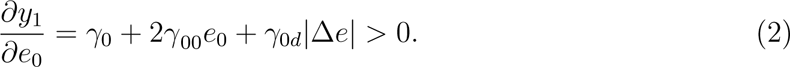

If the partial derivative of adult outcomes (*y*_1_) with respect to the quality of the developmental environment (*e*_0_) is not significantly positive (i.e., adult outcomes improve as a function of better developmental environments), then we cannot reject the null hypothesis that there is no relationship between the developmental environment and adult outcomes, contrary to the predictions of DC.

To test for AR following the definition given in Table 1, we use the following inequality:

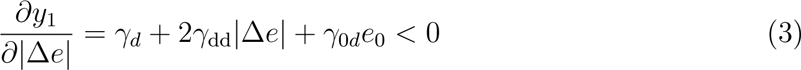

Here, if the partial derivative of adult outcomes (*y*_1_) with respect to the size of the *difference* between developmental and adult environments (*|*Δ*_e_|*) is not significantly negative (i.e., adult outcomes improve as a function of better matched developmental and adult environments), then we cannot reject the null that there is no relationship between the degree of environmental mismatch and adult outcomes, contrary to the predictions of AR. Further details on the theoretical motivation for the quadratic regression approach and the derivation of the associated hypothesis tests are available in [27]. To assist other authors with the implementation of quadratic models and the associated tests of early life environment and environmental delta derivatives, a link to R and Stata code written for this purpose can be found in the data availability statement.

Our models cluster on individual baboon IDs. We present the results in three ways: 1) with no fixed effects, which compares across individuals; 2) with a social group fixed effect, which compares across individuals within groups; and 3), for AR only, with a fixed effect corresponding to animal ID, which allows us to compare how changing levels of environmental mismatch are associated with fertility outcomes within individuals. We cannot test DC when including the individual-level fixed effect because each individual only experiences one developmental environment. The results are qualitatively similar across fixed-effects specifications, so we begin each section of the results with a general overview of the take-home message of the models. Any notable differences in effect sizes or significance levels are discussed in the text of the results section, and the full details of all three specifications are included in the results tables (Tables 4, 5, and 6).

**Table 4:** Results from quadratic models examining the effect of developmental environment and developmental/adult environment deltas on the probability of conceiving, given that a female was cycling at the start of the month.

|  | Rank |  |  | Rain |  |  |
| --- | --- | --- | --- | --- | --- | --- |
|  | (1) | (2) | (3) | (4) | (5) | (6) |
| $e_0$ | -0.179<br>(0.001) | -0.204<br>($<0.001$) | | -0.002<br>(0.319) | -0.002<br>(0.161) | |
| $\vert \Delta\vert = \vert e_1 - e_0\vert $ | -0.038<br>(0.634) | -0.031<br>(0.707) | 0.051<br>(0.728) | $<0.001$<br>(0.716) | 0.001<br>(0.430) | 0.001<br>(0.476) |
| $e_0^2$ | 0.171<br>(0.001) | 0.191<br>($<0.001$) | | $<0.001$<br>(0.164) | $<0.001$<br>(0.081) | |
| $e_0 \times \vert \Delta\vert $ | 0.049<br>(0.548) | 0.045<br>(0.613) | -0.027<br>(0.864) | $<-0.001$<br>(0.531) | $<-0.001$<br>(0.257) | $<-0.001$<br>(0.451) |
| $\vert \Delta\vert ^2$ | -0.021<br>(0.828) | -0.046<br>(0.643) | -0.054<br>(0.744) | $<0.001$<br>(0.775) | $<0.001$<br>(0.679) | $<0.001$<br>(0.799) |
| Age | 0.061<br>($<0.001$) | 0.064<br>($<0.001$) | 0.067<br>($<0.001$) | 0.060<br>($<0.001$) | 0.063<br>($<0.001$) | 0.067<br>($<0.001$) |
| Age sqd. | -0.002<br>($<0.001$) | -0.002<br>($<0.001$) | -0.002<br>($<0.001$) | -0.002<br>($<0.001$) | -0.002<br>($<0.001$) | -0.002<br>($<0.001$) |
| Group size | $<-0.001$<br>(0.428) | $<-0.001$<br>(0.244) | $<-0.001$<br>(0.702) | $<-0.001$<br>(0.381) | $<-0.001$<br>(0.322) | $<-0.001$<br>(0.638) |
| Observations | 8433 | 8433 | 8433 | 8404 | 8404 | 8404 |
| Sample means and sd: |  |  |  |  |  |  |
| $y$ | 0.137 | | | 0.137 | | |
|  | 0.344 |  |  | 0.344 |  |  |
| $e_0$ | 0.537 | | | 28.457 | | |
|  | 0.285 |  |  | 10.510 |  |  |
| $\vert \Delta\vert $ | 0.158 | | | 12.541 | | |
|  | 0.155 |  |  | 9.822 |  |  |
| Marginal effects and p: |  |  |  |  |  |  |
| DC: $\partial y / \partial e_0$ | 0.012<br>(0.383) | 0.009<br>(0.540) | | $<0.001$<br>(0.550) | $<-0.001$<br>(0.835) | |
| AR: $\partial y / \partial \vert \Delta\vert $ | -0.019<br>(0.617) | -0.021<br>(0.573) | 0.020<br>(0.749) | $<0.001$<br>(0.869) | $<0.001$<br>(0.643) | $<0.001$<br>(0.461) |
| Fixed effects | None | Group | Indiv. | None | Group | Indiv. |

**Table 5:**
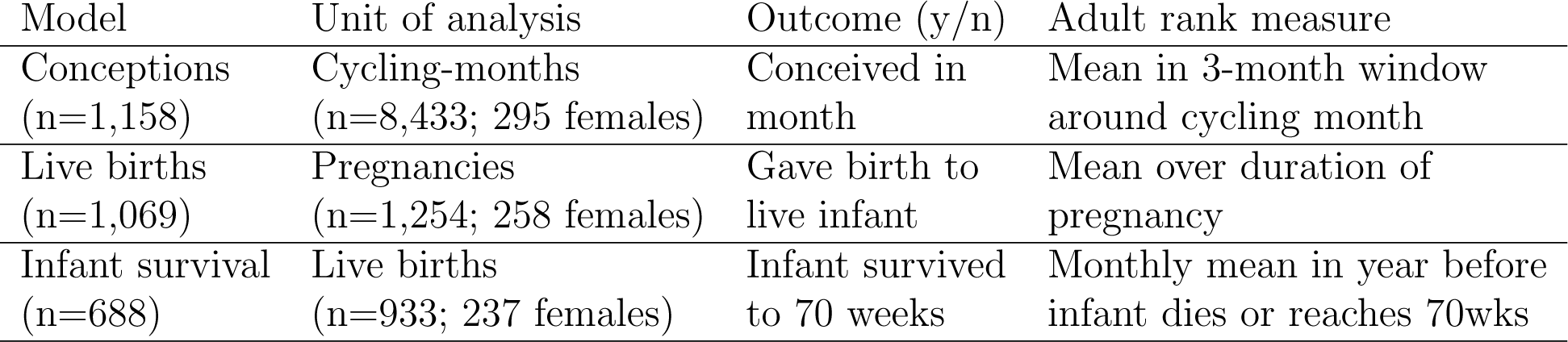
Results from quadratic models examining the effect of developmental environment and developmental/adult environment deltas on the probability of giving birth to a live infant, given that a female was pregnant.

Given the structure of the analyses (two hypotheses, two environmental variables of interest, with three fixed-effects specifications for each environmental variable), we inevitably conduct many hypothesis tests. We therefore provide both uncorrected p-values in the results tables and Bonferroni-corrected p-values in the text, where indicated [61, 62], as well as less conservative sharpened two-stage q-values that take into account the observed distribution of p-values across tests using a false discovery rate approach [63, 64].

## 3 Results

### 3.1 Tests of the developmental constraints hypothesis

#### 3.1.1 The effects of rainfall during development

None of our models provided any evidence that lower rainfall during development predicted worse fertility outcomes in adulthood. The amount of rainfall during development did not predict the likelihood of conceiving (contingent upon being cycling), giving birth to a live infant (contingent on pregnancy), or successfully raising an infant to 70 weeks (contingent on having given birth to a live infant). This was true regardless of whether we compared across all females (the models with no fixed effects, where *p >* 0.497 for all three outcomes; see column 4 in Tables 4, 5, and 6), or whether we compared females within groups (the group fixed effects models, where *p >* 0.523 for all three outcomes; see column 5 in Tables 4, 5, and 6).

#### 3.1.2 The effects of rank during development

Females who were lower-ranking during development were no less likely than their peers to conceive or to give birth to a live infant, regardless of whether the comparison was across females or within groups. In all cases, *p >* 0.382; see columns 1 and 2 in Tables 4 and 5.

Consistent with the developmental constraints hypothesis, within groups, females born to lower-ranking mothers were somewhat less likely to successfully raise an infant to 70 weeks than their peers who were born to higher-ranking mothers (*p* = 0.043; see column 2 in Table 6). In this model, females whose own mothers were in the 10th percentile of rank when they were born were 13.18% more likely to have their infant die before weaning than females whose mothers are in the 90th percentile of rank when they were born (SD=22.08%). Given that the mean infant survival probability was 73.74%, this translates to a 9.72% difference in the overall odds of infant survival for females in the 10th versus 90th rank percentiles (i.e., 13.18% of 73.74% is 9.72), or the equivalent of 25.55% of one standard deviation. The results were qualitatively similar when the comparison was across females (i.e., no fixed effects were included; *p* = 0.068, see column 1 in Table 6).

**Table 6:** Results from quadratic models examining the effect of developmental environment and developmental/adult environment deltas on the probability of raising an infant to 70 weeks, given that a female gave birth to a live infant.

|  | Rank |  |  | Rain |  |  |
| --- | --- | --- | --- | --- | --- | --- |
|  | (1) | (2) | (3) | (4) | (5) | (6) |
| $e_0$ | 0.649<br>(0.003) | 0.699<br>(0.002) | | 0.007<br>(0.252) | 0.006<br>(0.338) | |
| $\vert \Delta\vert = \vert e_1 - e_0\vert $ | -0.432<br>(0.146) | -0.369<br>(0.228) | -0.725<br>(0.232) | -0.002<br>(0.718) | 0.001<br>(0.843) | -0.001<br>(0.857) |
| $e_0^2$ | -0.542<br>(0.006) | -0.578<br>(0.005) | | <-0.001<br>(0.144) | <-0.001<br>(0.245) | |
| $e_0 \times \vert \Delta\vert $ | 0.102<br>(0.734) | 0.124<br>(0.682) | -0.208<br>(0.791) | <0.001<br>(0.247) | <0.001<br>(0.521) | <0.001<br>(0.478) |
| $\vert \Delta\vert ^2$ | 0.271<br>(0.475) | 0.298<br>(0.438) | 1.051<br>(0.144) | <-0.001<br>(0.803) | <-0.001<br>(0.560) | <-0.001<br>(0.893) |
| Age | 0.100<br>(<0.001) | 0.102<br>(<0.001) | 0.070<br>(0.005) | 0.105<br>(<0.001) | 0.108<br>(<0.001) | 0.070<br>(0.005) |
| Age sqd. | -0.004<br>(<0.001) | -0.004<br>(<0.001) | -0.003<br>(0.001) | -0.004<br>(<0.001) | -0.004<br>(<0.001) | -0.003<br>(<0.001) |
| Group size | 0.001<br>(0.517) | 0.001<br>(0.270) | 0.002<br>(0.165) | 0.001<br>(0.137) | 0.001<br>(0.234) | 0.002<br>(0.125) |
| Observations | 933 | 933 | 933 | 929 | 929 | 929 |
| Sample means and sd: |  |  |  |  |  |  |
| $y$ | 0.737 | | | 0.737 | | |
|  | 0.440 |  |  | 0.440 |  |  |
| $e_0$ | 0.529 | | | 29.049 | | |
|  | 0.299 |  |  | 11.217 |  |  |
| $\vert \Delta\vert $ | 0.167 | | | 14.066 | | |
|  | 0.161 |  |  | 11.347 |  |  |
| Marginal effects and p: |  |  |  |  |  |  |
| DC: $\partial y / \partial e_0$ | 0.093<br>(0.068) | 0.108<br>(0.043) | | <-0.001<br>(0.824) | <-0.001<br>(0.838) | |
| AR: $\partial y / \partial \vert \Delta\vert $ | -0.287<br>(0.038) | -0.204<br>(0.153) | -0.485<br>(0.081) | 0.001<br>(0.430) | 0.002<br>(0.354) | 0.001<br>(0.451) |
| Fixed effects | None | Group | Indiv. | None | Group | Indiv. |

Regardless of whether the comparison was within groups or across all females, the relationship between early life rank and infant survival was not statistically significant following adjustment for multiple tests (sharpened two-stage q-value = 0.690, while a Bonferroni-corrected threshold for statistical significance is *p <* 0.004).

### 3.2 Tests of the adaptive response hypothesis

#### 3.2.1 The effects of rain deltas

Females were not less likely to conceive, give birth to a live infant, or successfully raise an infant to 70 weeks when they experienced a large rainfall delta than when they experienced a small rainfall delta. Regardless of whether the comparison was across all females, within groups, or within individual females, *p >* 0.353 in all cases (see columns 4, 5, and 6 in Tables 4, 5, and 6).

#### 3.2.2 The effects of rank deltas

Females were not less likely to conceive or to give birth to a live infant when their dominance rank deltas were bigger than when they were smaller. Whether the comparison was across all females, within groups, or within individual females, *p >* 0.235 in all cases (see columns 1, 2, and 3 in Tables 4 and 5).

We identified no statistically significant patterns that supported the AR hypotheses, though there was one case in which results approached consistency with the mismatch prediction of AR. Comparing within individual animals, when females had larger dominance rank deltas they were more likely to lose an infant before weaning than when they had smaller rank deltas (where *p* = 0.081; individual fixed effects model in column 3 of Table 6; Figure 2). Based on a mean infant survival probability of 73.74%, this indicates that when a female was in the 90th percentile of rank difference she was 59.19% more likely to lose an infant before weaning than when she was in the 10th percentile of rank difference. This translates to a 43.65% difference in the overall odds of infant survival between the 10th and 90th percentiles of rank delta. Though this is a large effect size, we caution that the uncertainty in the effect size estimate is also high, with *p* values and sharpened q-values well above statistical significance thresholds after multiple testing adjustments. The results were qualitatively similar, though with smaller effect size estimates, when the comparison was across all females (59% of the within-individual effect size, unadjusted *p* = 0.038) or within groups (42% of the within-individual effect size, unadjusted *p* = 0.153; see columns 1 and 2 in Table 6).

**Figure 2:**
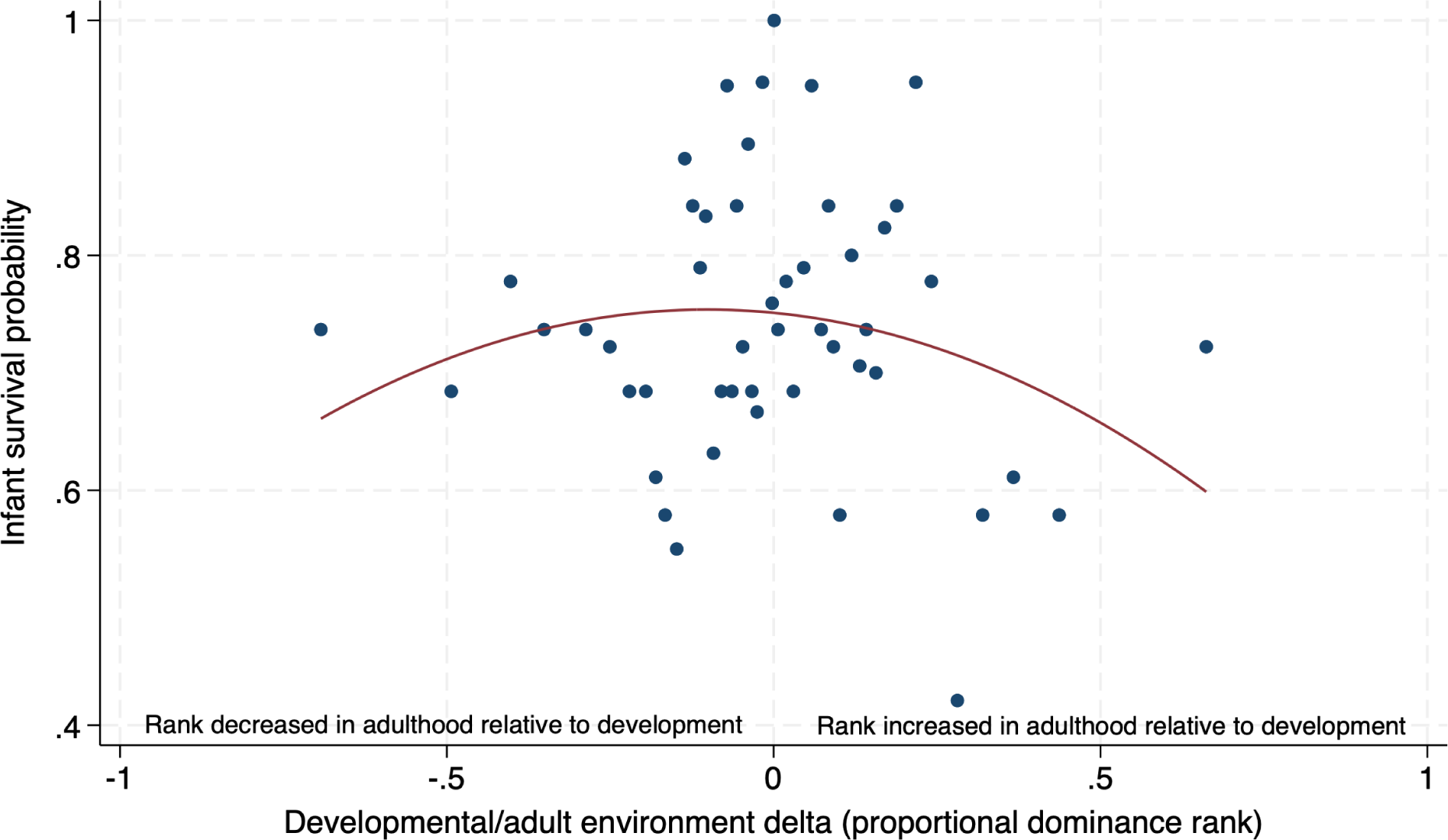
The relationship between dominance rank deltas and infant survival probability. Consistent with the predictions of the adaptive response (AR) hypothesis, we observed non-statistically significant evidence that females had a lower chance of conceiving when there were greater differences between their rank during development and their rank in adulthood than they did when these differences were smaller (unadjusted p-value of AR test on within-individual model=0.081, sharpened q-value=0.129; see column 3, Table 6). The center of the X axis (0) represents a developmental/adult environment delta of zero, meaning that the female held the same rank in adulthood as she did during development. Negative numbers represent periods during which a female fell in rank relative to the rank she experienced during development, while positive numbers represent periods during which she rose in rank relative to the rank she experienced during development. The plot shows raw (i.e. unadjusted) data grouped into 50 bins, overlaid with a quadratic fit line. Total sample size = 933 live births, so each bin contains *∼* 19 data points (2% of the sample).

## 4 Discussion

We find that low early-life rainfall and low early life social status–two important aspects of developmental environments for wild female baboons [4, 34, 41, 42]–do not play a substantive role in key female fertility outcomes for wild baboons. Being born during low-rainfall years or to a low-ranking mother did not significantly predict adult fertility parameters in our models, with one possible exception: females born to low-ranking mothers were somewhat more likely to have an infant die before weaning than females born to high-ranking mothers were. However, this effect is not statistically well-supported in our sample, which is large compared to many studies of wild primates.

Our results therefore do not provide clear evidence in favor of either DC or AR for the developmental environment variables and adult outcomes we considered here. This deviates from a prior analysis done on this baboon population [34]. That study concluded that drought conditions early in life predicted reduced probabilities of conception and resumption of cycling following post-partum amenorrhea (relative to females who did not experience drought) specifically when females experienced drought conditions in adulthood [34]. We interpreted these results as support for DC because females born in drought conditions did worse when faced with drought again in adulthood than they did when their adult environments were more favorable. The earlier analysis differs substantially from the present analysis in the way the dataset was constructed: the previous analysis focused on females born in different early life rainfall environments (treated as a binary variable) and asked whether fertility outcomes differed for females that then lived through both a normal rainfall year and an extreme drought in adulthood (n=50 females fit these criteria). Some of the differences in findings are thus likely related to differences in the composition and size of the datasets. Others are likely an outgrowth of analysis strategy: [34] controlled for reproductive state rather than conditioning females’ inclusion in a given analysis on reproductive state. Furthermore, the earlier analysis relied on a model containing an interaction term between the developmental and adult environments. Here, we instead used a quadratic regression and evaluated the partial derivatives of outcomes with respect to the quality of the developmental environment (for DC) and the size of the difference between early and adult environments (for AR). These are analysis decisions that we have previously argued increase the reliability and interpretability of tests of the DC and AR hypotheses [27].

Our findings emphasize the importance of methodological choices when testing these hypotheses, which may be one of several potential reasons why the literature is still unclear on their relative importance in humans and other animals. While prior literature has generally found more support for the DC hypothesis than for AR hypotheses [reviewed in 14, 16, 25], there is considerable heterogeneity across the literature. For example, in support of DC, better early life environments are associated with greater lifetime reproductive success in bighorn sheep, Svalbard reindeer, and spotted hyenas [23, 65, 66]. Meanwhile, three recent studies of wild mammals have not found clear evidence for decreased fitness in response to some kinds of early life adversity. African elephants who lose their herd matriarch (not their mother) early in life do not appear to suffer fitness costs, nor do female mountain gorillas who lose their mother at a relatively young age [67, 68]. Furthermore, drought in early life is not associated with reduced fitness in female elephants or banded mongooses [68, 69].

Mammalian studies with support for the environmental mismatch prediction of AR hypotheses are scarce. However, some have found evidence for AR in the form of phenotypes that develop as a result of exposure to certain environments *in utero* or very early life. That is, instead of testing whether environmental deltas predict adult outcomes–the strategy we used here–they tested whether developmental environments predicted the appearance of a specific phenotype, a complementary and non-mutually-exclusive testing strategy [27]. For example, pregnant voles exposed to light cues which signal that pups will be born during the winter gave birth to pups that were born with thicker coats. In contrast, the pups of females who received light cues indicating a warm birth environment were born with thinner coats [70]. Red squirrel pups exposed to cues (including *in utero*) indicating they lived in a high population density environment grew faster than pups who did not receive such cues, and faster growth increased the likelihood of survival to potential reproductive age specifically in these high-density environments [71].

One of the challenges in resolving AR and DC explanations is fundamental to any study of a natural, un-manipulated population: it is impossible to vary only the predictors of interest while holding all else constant. Consequently, researchers must assume that the predictor of interest is randomly distributed across study subjects. In our study, this is probably a safe assumption in the case of rainfall, but less clear in the case of dominance rank. Females who hold different ranks during development are likely different from one another in a variety of ways that affect later life fertility, such as growth rates, adult size, or resource access [41, 57, 72]. And because female baboons non-genetically “inherit” their rank from their mothers (e.g., in our infant survival data set, the correlation between *e*0 and *e*1 is 0.71) [43, 44], rank-related DC may simply arise from an effect of rank in adulthood. These intrinsic design issues complicate the interpretation of results, making experimental work on this subject especially valuable [71, 73].

For the AR hypothesis, our data allowed us to perform within-individual comparisons across adult years, somewhat mitigating the problem of confounding between-animal differences. However, even here, there are a variety of ways in which the experiences and environments of female baboons might be different when their rank deltas are large versus when they are small. For example, rising in rank is disproportionately associated with group fissions. Fissions are most likely to occur when animals are experiencing resource stress associated with large group size [74, 75] and disrupts the lives of females in the study population, even if they mean an increase in status. Meanwhile, falling in rank may occur because females are sick or injured, making them vulnerable to displacement by lower-ranking females. Consequently, multiple pathways can give rise to environmental mismatch effects, some of which also can independently generate poor outcomes without requiring a causal effect of phenotypic “choices” organisms make. For example, in wild roe deer, increased viability selection in poor-quality environments may generate apparent fitness benefits to females who experienced “matched” poor-quality developmental and adult environments: because high-quality females were more likely to survive the harsh early environment, they performed better than the (on average) lower-quality females born in favorable environments, when both groups encountered harsh conditions later in life [76]. Most studies of human populations have ignored the potential role of viability selection and other alternative explanations that may generate a connection between environmental deltas and adult outcomes [e.g., 30]. Finally, the relative lack of evidence we find for effects of early life adversity on fertility measures suggests that shortened lifespans are the primary mechanism by which early adversity might decrease lifetime fitness. Prior analyses have demonstrated that experiencing more sources of early life adversity, including drought and being born to a low-ranking mother, leads to markedly shorter lifespans in this baboon population [4, 37, 77]. Lifespan is by far the single biggest contributor to females’ lifetime reproductive success, explaining 80-90% of the observed variation in the Amboseli population [57, 77]. Indeed, across taxa, shortened lifespans appear to be a common consequence of early life adversity, accounting for a significant proportion of the studies that provide evidence for DC (e.g. red squirrels: [73]; chimpanzees: [78]; Asian elephants: [79]; hyenas: [80]; reviewed in [26, 39]). Consequently, for long-lived species like primates, lifespan analyses will likely be crucial to understanding the extent to which early life adversity compromises fitness.

In sum, our data do not provide clear support for either the developmental constraints or adaptive response hypothesis as compelling explanations for differences in fertility outcomes in wild female baboons, at least when social status and rainfall are the environmental variables of interest. We hope that these analyses, which are structured to help avoid empirical conflation of the hypotheses, will motivate additional evaluation of the evidence for DC and AR using the statistical methods demonstrated here.

## 5 Data availability statement

The data reported in this paper can be found in the Dryad repository at https://doi.org.10.5061/dryad.2v6wwpzw9. The code needed to replicate the analysis is available on GitHub at https://github.com/anup-malani/rosenbaum_etal_2024_baboon_dc_ar_mismatch. To assist other authors with the implementation of quadratic models and the associated tests of early life environment and environmental delta derivatives, a link to R and Stata code written for this purpose can be found at https://github.com/anup-malani/PAR.

## Acknowledgements

We gratefully acknowledge the support of the National Science Foundation and the National Institutes of Health for the majority of the data represented here, currently through R01AG071684, R01AG075914, and R61AG078470. Current support for field-based data collection also comes from the Max Planck Institute for Evolutionary Anthropology, and we thank Duke University, Princeton University, and the University of Notre Dame for financial and logistical support over the years. In Kenya, our research was approved by the Wildlife Research Training Institute (WRTI), Kenya Wildlife Service (KWS), the National Commission for Science, Technology, and Innovation (NACOSTI), and the National Environment Management Authority (NEMA). We also thank the University of Nairobi, the Institute of Primate Research (IPR), the National Museums of Kenya, the members of the Amboseli-Longido pastoralist communities, the Enduimet Wildlife Management Area, Ker & Downey Safaris, Air Kenya, and Safarilink for their cooperation and assistance in the field. Particular thanks go to the Amboseli Baboon Project long-term field team (R.S. Mututua, S. Sayialel, J.K. Warutere, I.L. Siodi, I.L., and L. Musembei), and to T. Wango and V. Oudu for their untiring assistance in Nairobi. The baboon project database, Babase, is expertly managed by N. Learn, J. Gordon, and W. Wilbur. Database design and programming are provided by K. Pinc. This research was approved by the IACUC at Duke University, University of Notre Dame, and Princeton University and the Ethics Council of the Max Planck Society and adhered to all the laws and guidelines of Kenya. For a complete set of acknowledgments of funding sources, logistical assistance, and data collection and management, please visit http://amboselibaboons.nd.edu/acknowledgements/. AM acknowledges the support of the Barbara J. and B. Mark Fried Fund at the University of Chicago Law School.

## Supplementary Materials

### A Rainfall and rank summary statistics

The following three tables contain the sample sizes and summary statistics for rainfall and dominance rank, the two measures we used to evaluate the developmental constraints hypothesis and the adaptive response hypothesis. *e*_0_ indicates rain (or rank) during development, while *e*_1_ indicates rain (or rank) in adulthood. Sample sizes differ between the two measures because *e*_0_ rainfall data were not available for the first two years included in our dataset (1974-75). See also Figure 1 in the main text, which shows the distributions of the difference between *e*_0_ and *e*_1_ rainfall and rank for each of our three data sets (where conceptions, live births, and infant survival are the outcome variables, presented respectively from left to right across the figure).

**Table A.1:** Rainfall (mm/month) and proportional dominance rank summary statistics for conception model.

| Variable | Sample size | Mean | Std Dev | Min | Max |
| --- | --- | --- | --- | --- | --- |
| Rain ($e_0$) | 8404 | 28.46 | 10.51 | 7.64 | 63.92 |
| Rain ($e_1$) | 8404 | 29.52 | 12.03 | 9.13 | 65.03 |
| Rank ($e_0$) | 8433 | 0.54 | 0.28 | 0.00 | 1.00 |
| Rank ($e_1$) | 8433 | 0.51 | 0.30 | 0.00 | 1.00 |

**Table A.2:**
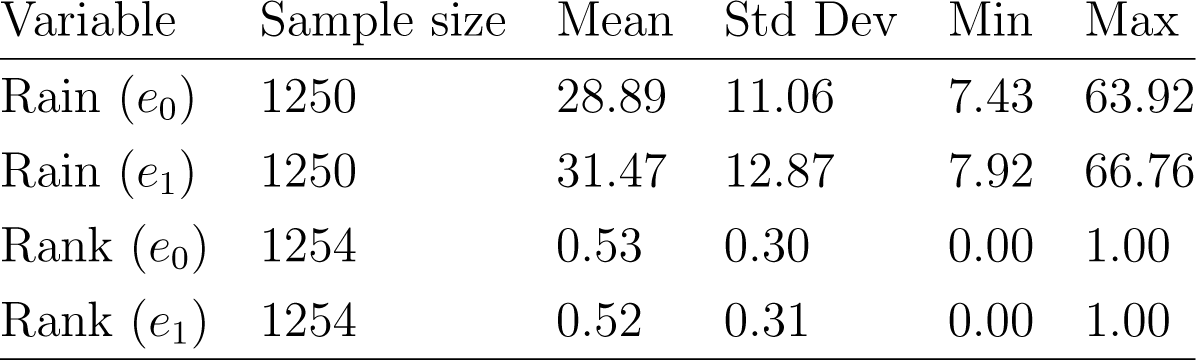
Rainfall (mm/month) and proportional rank summary statistics for live birth model.

**Table A.3:**
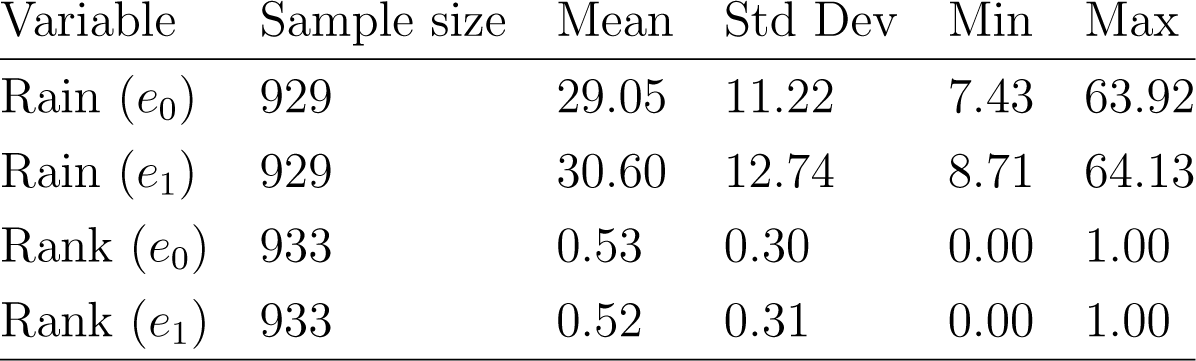
Rainfall (mm/month) and proportional rank summary statistics for infant survival model.

| Variable | Sample size | Mean | Std Dev | Min | Max |
| --- | --- | --- | --- | --- | --- |
| Rain ($e_0$) | 929 | 29.05 | 11.22 | 7.43 | 63.92 |
| Rain ($e_1$) | 929 | 30.60 | 12.74 | 8.71 | 64.13 |
| Rank ($e_0$) | 933 | 0.53 | 0.30 | 0.00 | 1.00 |
| Rank ($e_1$) | 933 | 0.52 | 0.31 | 0.00 | 1.00 |

### B Dominance rank data

Female baboons form strong, stable dominance hierarchies that determine access to important resources [52]. Female infants “inherit” their ranks from their mothers via a system of youngest ascendancy, where a female’s youngest daughter becomes dominant over her older maternal sisters when she attains adult rank (typically between 2 and 3 years old, and often with the assistance of her mother). This means that female baboons can reasonably predict that they will hold a dominance rank similar to the one their mother held when they were born, when they themselves are adults. However, events such as group fissions and matriline overthrows [32] mean that some individuals will hold dominance ranks in adulthood that are either higher or lower than the dominance rank their mother held when they were born. Since there is variation in both developmental environment (some baboons are born to low-ranking mothers, and some to high-ranking ones) and in how well the developmental and adult environments match (some females will hold nearly the same rank their whole lives, and some will experience a great deal of change), rank is an appropriate variable for testing both the developmental constraints and adaptive response hypotheses.

We used proportional (i.e., relative) dominance ranks, which represent the proportion of adult female group members that the subject in question outranks [51, 52]. Each animal’s rank was determined based on the outcomes of all observed, decided agonistic interactions between adult females [53]. Trained observers, who recognize individual animals via differences in morphological characteristics, record the identities of individuals participating in agonistic encounters and the outcome of each of these encounters. If one animal behaved submissively while the other was either aggressive or remained neutral, the interaction was recorded as “decided” with respect to the rank relationship between the two interacting animals. Any interactions without a clear outcome (e.g., where both animals displayed submissive signals) were excluded from dominance rank calculations. We eliminated periods of rank instability (e.g., when groups were going through fissions, during which rank can be difficult to determine).

### C Model covariates

Variables besides those of interest here may influence female fertility, in particular age and group size [34, 40, 54]. We included variables known to be important from previous analyses as covariates in our models. Group size data came from group censuses conducted during routine research visits; females ages were known from long-term monitoring of the population. Covariate definitions and descriptive statistics are shown in Table A.4.

**Table A.4:**
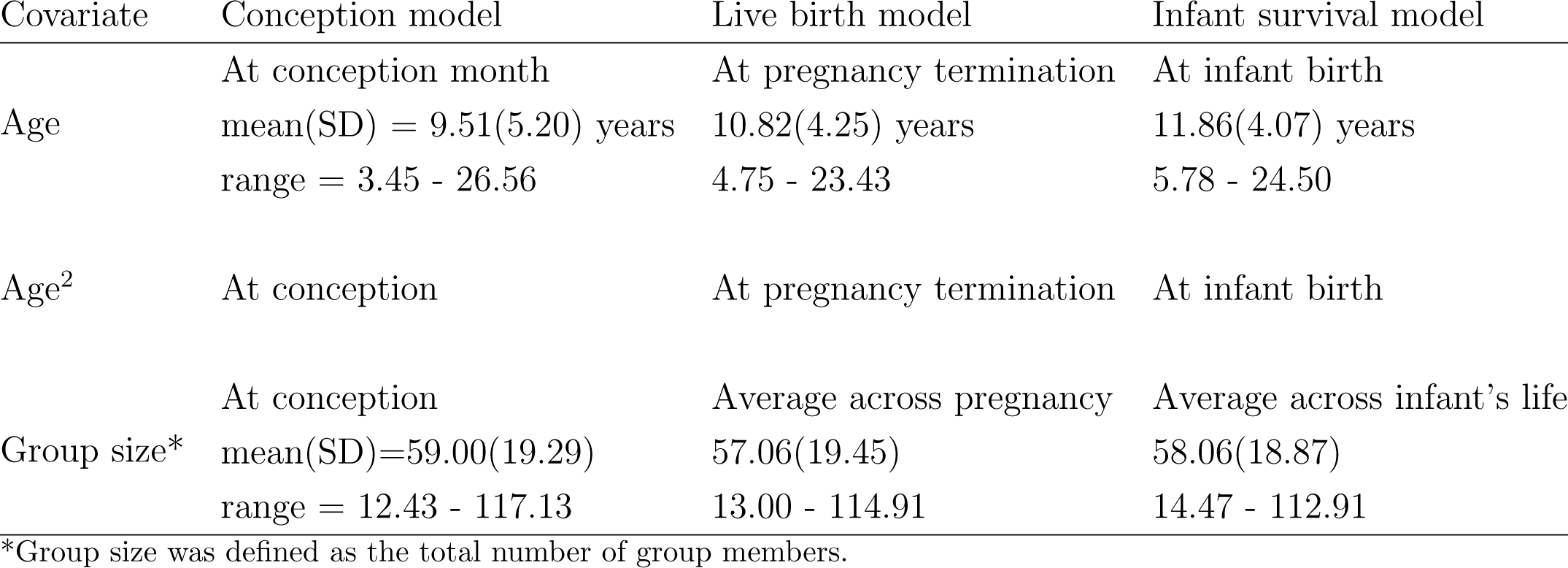
Description and summary statistics of covariates used in regression models evaluating developmental constraints and adaptive responses in female baboons.

### D Conception models with reduced sample

The below table (Table A.5) presents results from quadratic models examining the effect of dominance rank (left three columns) and rainfall (right three columns) on the probability of successfully conceiving given that 1) the female was cycling at the start of the month, and 2) the female’s last pregnancy resulted in a live infant who lived to at least 70 weeks. That is, the data set used to generate this table excludes nulliparous females and females whose prior infant died before weaning age, both of which are known from prior research to require more cycles before conceiving. These results can be compared to the models in Table 4 in the main text; Table 4 is structured identically but uses a data set that also contains data from nulliparous females and females whose prior pregnancy resulted in the loss of an infant.

Unlike for the conception results in presented in the main text, in this reduced sample females were less likely to conceive when the differences between their developmental rank and adult rank were bigger in two out of the three models (where *p* = 0.055 and *p* = 0.071). One of these models makes a between-animal comparison, i.e., it does not contain any fixed effects (column 1, where *p* = 0.055), and the other makes a within-group comparison, i.e., it contains a social group fixed effect (column 2, where *p* = 0.071). In this subsample, females in the 10th percentile of developmental rank/adult rank difference (i.e., their ranks were similar) were 63% (SD=30.61%) more likely to conceive than females in the 90th percentile of rank difference. Note that neither of these results comes close to surviving any version of a multiple testing adjustment. When the comparison was conducted within individual females this result was not statistically significant even without a multiple testing adjustment (column 3, individual fixed effects, *p* = 0.946). As we have argued in the main text, this is the most powerful of the three tests of the effects of developmental and adult environmental mismatch.

**Table A.5:**
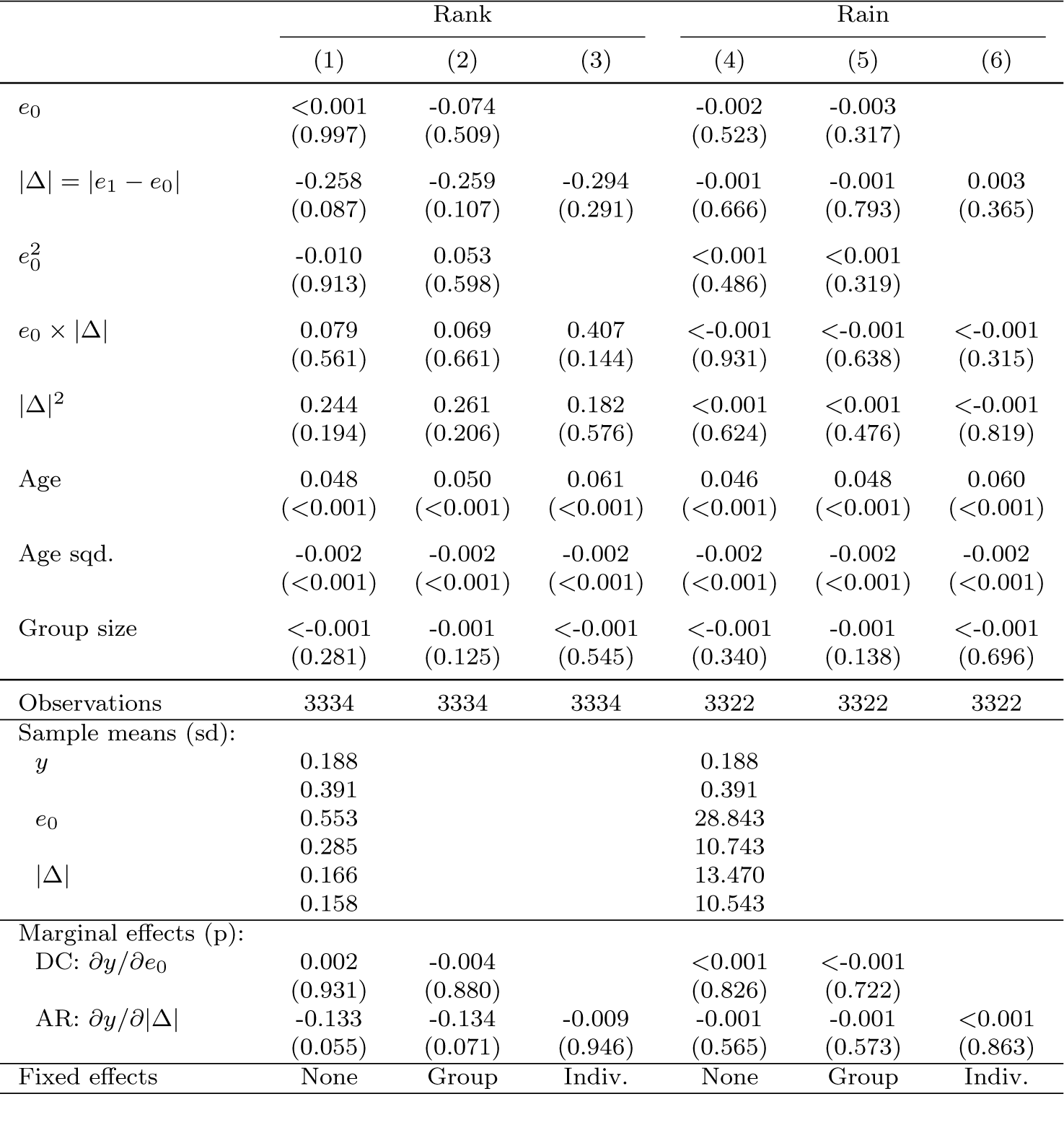
Results from quadratic models examining the effect of developmental environment and developmental/adult environment deltas on the probability of conceiving, given that a female was cycling at the start of the month.

Results from quadratic models examining the effect of dominance rank and rainfall on the probability of conceiving, given that a female was cycling at the start of a given observation month. Rank results can be found in columns 1-3, which correspond to models with no fixed effect, a group-level fixed effect, and an individual-level fixed-effect respectively; rainfall results are in columns 4-6, with the same three fixed effects respectively. In the first panel, each cell provides the coefficient associated with a given model term, and below it, the p-value in parentheses. In the second panel, each cell provides means with standard deviations below it. In the third panel, each cell provides marginal effects at the mean value of independent variables, and the p-value in parentheses below it. *y* is the dependent variable, i.e., outcome. *e*0 is developmental environment and Δe is the difference between developmental and adult environment.

## References

[1] David JP Barker and Clive Osmond. Infant mortality, childhood nutrition, and ischaemic heart disease in England and Wales. The Lancet, 327(8489):1077–1081, 1986. ISSN 0140-6736. URL https://www.sciencedirect.com/science/article/abs/pii/S0140673686913401.

[2] Vincent J Felitti, Robert F Anda, Dale Nordenberg, David F Williamson, Alison M Spitz, Valerie Edwards, and James S Marks. Relationship of childhood abuse and household dysfunction to many of the leading causes of death in adults: The adverse childhood experiences (ACE) study. American Journal of Preventive Medicine, 14(4): 245–258, 1998. ISSN 0749-3797. URL https://www.sciencedirect.com/science/article/abs/pii/S0749379798000178.

[3] Peter D Gluckman, Mark A Hanson, and Alan S Beedle. Early life events and their consequences for later disease: a life history and evolutionary perspective. American Journal of Human Biology, 19(1):1–19, 2007. ISSN 1042-0533. URL https://onlinelibrary.wiley.com/doi/abs/10.1002/ajhb.20590.

[4] Jenny Tung, Elizabeth A. Archie, Jeanne Altmann, and Susan C. Alberts. Cumulative early life adversity predicts longevity in wild baboons. Nature Communications, 7(1): 11181, 2016. ISSN 2041-1723. doi: 10.1038/ncomms11181. URL 10.1038/ncomms11181.

[5] Andreas Berghanel, Michael Heistermann, Oliver Schülke, and Julia Ostner. Prenatal stress effects in a wild, long-lived primate: predictive adaptive responses in an unpredictable environment. Proceedings of the Royal Society B: Biological Sciences, 283(1839):20161304, 2016. doi: 10.1098/rspb.2016.1304. URL https://royalsocietypublishing.org/doi/abs/10.1098/rspb.2016.1304.

[6] Harrison J.F. Eyck, Katherine L. Buchanan, Ondi L. Crino, and Tim S. Jessop. Effects of developmental stress on animal phenotype and performance: a quantitative review. Biological Reviews, 94(3):1143–1160, 2019. ISSN 1464-7931. doi: 10.1111/brv.12496. URL https://onlinelibrary.wiley.com/doi/abs/10.1111/brv.12496.

[7] Sam K. Patterson, Katie Hinde, Angela B. Bond, Benjamin C. Trumble, Shirley C. Strum, and Joan B. Silk. Effects of early life adversity on maternal effort and glucocorticoids in wild olive baboons. Behavioral Ecology and Sociobiology, 75(8):114, 2021. ISSN 1432-0762. doi: 10.1007/s00265-021-03056-7. URL 10.1007/s00265-021-03056-7.

[8] Jan Lindstrom. Early development and fitness in birds and mammals. Trends in Ecology & Evolution, 14(9):343–348, 1999. ISSN 0169-5347. doi: 10.1016/S0169-5347(99)01639-0. URL http://www.sciencedirect.com/science/article/pii/S0169534799016390.

[9] Peter D Gluckman and Mark A Hanson. Developmental origins of disease paradigm: a mechanistic and evolutionary perspective. Pediatric Research, 56(3):311–317, 2004. ISSN 1530-0447. URL https://www.nature.com/articles/pr2004210.

[10] Jay Belsky and Michael Pluess. Beyond diathesis stress: differential susceptibility to environmental influences. Psychological Bulletin, 135(6):885, 2009. ISSN 1939-1455. URL https://psycnet.apa.org/record/2009-19763-005.

[11] Christopher W. Kuzawa and Elizabeth A. Quinn. Developmental origins of adult function and health: Evolutionary hypotheses. Annual Review of Anthropology, 38 (1):131–147, 2009. doi: 10.1146/annurev-anthro-091908-164350. URL https://www.annualreviews.org/doi/abs/10.1146/annurev-anthro-091908-164350.

[12] Daniel Nettle, Willem E. Frankenhuis, and Ian J. Rickard. The evolution of predictive adaptive responses in human life history. Proceedings of the Royal Society B: Biological Sciences, 280(1766):20131343, 2013. doi: 10.1098/rspb.2013.1343. URL https://royalsocietypublishing.org/doi/abs/10.1098/rspb.2013.1343.

[13] Carlos A Botero, Franz J Weissing, Jonathan Wright, and Dustin R Rubenstein. Evolutionary tipping points in the capacity to adapt to environmental change. Proceedings of the National Academy of Sciences, 112(1):184–189, 2015. ISSN 0027-8424. URL https://www.pnas.org/doi/abs/10.1073/pnas.1408589111.

[14] Amy Lu, Lauren Petrullo, Sofia Carrera, Jacob Feder, India Schneider-Crease, and Noah Snyder-Mackler. Developmental responses to early-life adversity: Evolutionary and mechanistic perspectives. Evolutionary Anthropology, 28(5):249–266, 2019. ISSN 1060-1538. doi: 10.1002/evan.21791. URL https://onlinelibrary.wiley.com/doi/abs/10.1002/evan.21791.

[15] Pat Monaghan. Early growth conditions, phenotypic development and environmental change. Philosophical Transactions of the Royal Society B: Biological Sciences, 363(1497):1635–1645, 2008. doi: 10.1098/rstb.2007.0011. URL https://royalsocietypublishing.org/doi/abs/10.1098/rstb.2007.0011.

[16] Amanda J. Lea and Stacy Rosenbaum. Understanding how early life effects evolve: progress, gaps, and future directions. Current Opinion in Behavioral Sciences, 36: 29–35, 2020. ISSN 2352-1546. doi: 10.1016/j.cobeha.2020.06.006. URL http://www.sciencedirect.com/science/article/pii/S2352154620300942.

[17] Peter D. Gluckman, Mark A. Hanson, and Felicia M. Low. Evolutionary and developmental mismatches are consequences of adaptive developmental plasticity in humans and have implications for later disease risk. Philosophical Transactions of the Royal Society B: Biological Sciences, 374(1770):20180109, 2019. doi: 10.1098/rstb.2018.0109. URL https://royalsocietypublishing.org/doi/abs/10.1098/rstb.2018.0109.

[18] Peter D Gluckman, Mark A Hanson, Hamish G Spencer, and Patrick Bateson. Environmental influences during development and their later consequences for health and disease: implications for the interpretation of empirical studies. Proceedings of the Royal Society B: Biological Sciences, 272(1564):671–677, 2005. doi: doi:10.1098/rspb.2004. 3001. URL https://royalsocietypublishing.org/doi/abs/10.1098/rspb.2004.3001.

[19] Adam D. Hayward and Virpi Lummaa. Testing the evolutionary basis of the predictive adaptive response hypothesis in a preindustrial human population. Evolution, Medicine, and Public Health, 2013(1):106–117, 2013. ISSN 2050-6201. doi: 10.1093/emph/eot007. URL 10.1093/emph/eot007.

[20] Daniel Nettle and Melissa Bateson. Adaptive developmental plasticity: what is it, how can we recognize it and when can it evolve? Proceedings of the Royal Society B: Biological Sciences, 282(1812):20151005, 2015. doi: 10.1098/rspb.2015.1005. URL https://royalsocietypublishing.org/doi/abs/10.1098/rspb.2015.1005.

[21] Andreas Berghänel, Michael Heistermann, Oliver Schülke, and Julia Ostner. Prenatal stress accelerates offspring growth to compensate for reduced maternal investment across mammals. Proceedings of the National Academy of Sciences, 114(50):E10658– E10666, 2017. ISSN 0027-8424. URL https://www.pnas.org/doi/abs/10.1073/pnas.1707152114.

[22] Milind Watve. Developmental plasticity: Need to go beyond näıve thinking. Evolution, Medicine, and Public Health, 2017(1):178–180, 2018. ISSN 2050-6201. doi: 10.1093/emph/eox020. URL 10.1093/emph/eox020.

[23] Gabriel Pigeon, Leif Egil Loe, Richard Bischof, Christophe Bonenfant, Mads Forchhammer, R. Justin Irvine, Erik Ropstad, Audun Stien, Vebjørn Veiberg, and Steve Albon. Silver spoon effects are constrained under extreme adult environmental conditions. Ecology, 100(12):e02886, 2019. ISSN 0012-9658. doi: 10.1002/ecy.2886. URL https://esajournals.onlinelibrary.wiley.com/doi/abs/10.1002/ecy.2886.

[24] Valeria Marasco, Steve Smith, and Fŕederic Angelier. How does early-life adversity shape telomere dynamics during adulthood? problems and paradigms. BioEssays, 44 (4):2100184, 2022. ISSN 0265-9247. URL https://onlinelibrary.wiley.com/doi/full/10.1002/bies.202100184.

[25] Amanda J Lea, Jenny Tung, Elizabeth A Archie, and Susan C Alberts. Developmental plasticity: Bridging research in evolution and human health. Evolution, Medicine, and Public Health, 2017(1):162–175, 2018. ISSN 2050-6201. doi: 10.1093/emph/eox019. URL 10.1093/emph/eox019.

[26] Amanda M. Dettmer and Daniella E. Chusyd. Early life adversities and lifelong health outcomes: A review of the literature on large, social, long-lived nonhuman mammals. Neuroscience & Biobehavioral Reviews, 152:105297, 2023. ISSN 0149-7634. doi: 10.1016/j.neubiorev.2023.105297. URL https://www.sciencedirect.com/science/article/pii/S014976342300266X.

[27] Anup Malani, Elizabeth A. Archie, and Stacy Rosenbaum. Conceptual and analytical approaches for modelling the developmental origins of inequality. Philosophical Transactions of the Royal Society B: Biological Sciences, 378(1883):20220306, 2023. doi: 10.1098/rstb.2022.0306. URL https://royalsocietypublishing.org/doi/abs/10.1098/rstb.2022.0306.

[28] Patrick Bateson. Fetal experience and good adult design. International Journal of Epidemiology, 30(5):928–934, 2001. ISSN 0300-5771. doi: 10.1093/ije/30.5.928. URL 10.1093/ije/30.5.928.

[29] Ian J. Rickard and Virpi Lummaa. The predictive adaptive response and metabolic syndrome: challenges for the hypothesis. Trends in Endocrinology & Metabolism, 18 (3):94–99, 2007. ISSN 1043-2760. doi: 10.1016/j.tem.2007.02.004. URL https://www.sciencedirect.com/science/article/pii/S1043276007000094.

[30] Jonathan CK Wells. A critical appraisal of the predictive adaptive response hypothesis. International Journal of Epidemiology, 41(1):229–235, 2012. ISSN 1464-3685. URL https://academic.oup.com/ije/article/41/1/229/651311.

[31] Patrick Bateson, Peter Gluckman, and Mark Hanson. The biology of developmental plasticity and the predictive adaptive response hypothesis. The Journal of Physiology, 592(11):2357–2368, 2014. ISSN 0022-3751. doi: 10.1113/jphysiol. 2014.271460. URL https://physoc.onlinelibrary.wiley.com/doi/abs/10.1113/jphysiol.2014.271460.

[32] Susan C. Alberts and Jeanne Altmann. The Amboseli Baboon Research Project: 40 years of continuity and change, pages 261–287. Springer Berlin Heidelberg, Berlin, Heidelberg, 2012. ISBN 978-3-642-22514-7. doi: 10.1007/978-3-642-22514-712. URL 10.1007/978-3-642-22514-7_12.

[33] Sam K. Patterson, Shirley C. Strum, and Joan B. Silk. Early life adversity has long-term effects on sociality and interaction style in female baboons. Proceedings of the Royal Society B: Biological Sciences, 289(1968):20212244, 2022. doi: 10.1098/rspb.2021.2244. URL https://royalsocietypublishing.org/doi/abs/10.1098/rspb.2021.2244.

[34] Amanda J. Lea, Jeanne Altmann, Susan C. Alberts, and Jenny Tung. Developmental constraints in a wild primate. The American Naturalist, 185(6):809–821, 2015. ISSN 0003-0147. doi: 10.1086/681016. URL 10.1086/681016.

[35] Matthew N Zipple, Elizabeth A Archie, Jenny Tung, Jeanne Altmann, and Susan C Alberts. Intergenerational effects of early adversity on survival in wild baboons. Elife, 8: e47433, 2019. ISSN 2050-084X. URL https://elifesciences.org/articles/47433.

[36] Elizabeth C. Lange, Shuxi Zeng, Fernando A. Campos, Fan Li, Jenny Tung, Elizabeth A. Archie, and Susan C. Alberts. Early life adversity and adult social relationships have independent effects on survival in a wild primate. Science Advances, 9(20):eade7172, 2023. doi: 10.1126/sciadv.ade7172. URL https://www.science.org/doi/abs/10.1126/sciadv.ade7172.

[37] Jenny Tung, Elizabeth C. Lange, Susan C. Alberts, and Elizabeth A. Archie. Social and early life determinants of survival from cradle to grave: A case study in wild baboons. Neuroscience & Biobehavioral Reviews, 152:105282, 2023. ISSN 0149-7634. doi: 10.1016/j.neubiorev.2023.105282. URL https://www.sciencedirect.com/science/article/pii/S0149763423002518.

[38] Willem E. Frankenhuis, Daniel Nettle, and Sasha R. X. Dall. A case for environmental statistics of early-life effects. Philosophical Transactions of the Royal Society B: Biological Sciences, 374(1770):20180110, 2019. doi: 10.1098/rstb.2018.0110. URL https://royalsocietypublishing.org/doi/abs/10.1098/rstb.2018.0110.

[39] Noah Snyder-Mackler, Joseph Robert Burger, Lauren Gaydosh, Daniel W. Belsky, Grace A. Noppert, Fernando A. Campos, Alessandro Bartolomucci, Yang Claire Yang, Allison E. Aiello, Angela O’Rand, Kathleen Mullan Harris, Carol A. Shively, Susan C. Alberts, and Jenny Tung. Social determinants of health and survival in humans and other animals. Science, 368(6493):eaax9553, 2020. doi: 10.1126/science.aax9553. URL https://science.sciencemag.org/content/sci/368/6493/eaax9553.full.pdf.

[40] Jacinta C. Beehner, Daphne A. Onderdonk, Susan C. Alberts, and Jeanne Altmann. The ecology of conception and pregnancy failure in wild baboons. Behavioral Ecology, 17(5):741–750, 2006. ISSN 1045-2249. doi: 10.1093/beheco/arl006. URL 10.1093/beheco/arl006.

[41] Susan C. Alberts. Social influences on survival and reproduction: Insights from a long-term study of wild baboons. Journal of Animal Ecology, 88(1):47–66, 2019. ISSN 0021-8790. doi: 10.1111/1365-2656.12887. URL https://besjournals.onlinelibrary.wiley.com/doi/abs/10.1111/1365-2656.12887.

[42] Emily J. Levy, Anna Lee, I. Long’ida Siodi, Emma C. Helmich, Emily M. McLean, Elise J. Malone, Maggie J. Pickard, Riddhi Ranjithkumar, Jenny Tung, Elizabeth A. Archie, and Susan C. Alberts. Early life drought predicts components of adult body size in wild female baboons. American Journal of Biological Anthropology, 182(3):357– 371, 2023. ISSN 2692-7691. doi: 10.1002/ajpa.24849. URL https://onlinelibrary.wiley.com/doi/abs/10.1002/ajpa.24849.

[43] Glenn Hausfater, Jeanne Altmann, and Stuart Altmann. Long-term consistency of dominance relations among female baboons (Papio cynocephalus). Science, 217(4561): 752–755, 1982. ISSN 0036-8075. URL https://www.science.org/doi/abs/10.1126/science.217.4561.752.

[44] Amanda J. Lea, Niki H. Learn, Marcus J. Theus, Jeanne Altmann, and Susan C. Alberts. Complex sources of variance in female dominance rank in a nepotistic society. Animal Behaviour, 94:87–99, 2014. ISSN 0003-3472. doi: 10.1016/j.anbehav.2014.05.019. URL https://www.sciencedirect.com/science/article/pii/S0003347214002395.

[45] J. Altmann, S. C. Alberts, S. A. Altmann, and S. B. Roy. Dramatic change in local climate patterns in the Amboseli basin, Kenya. African Journal of Ecology, 40(3): 248–251, 2002. ISSN 0141-6707. doi: 10.1046/j.1365-2028.2002.00366.x. URL https://onlinelibrary.wiley.com/doi/abs/10.1046/j.1365-2028.2002.00366.x.

[46] Mildred M Aduma, Gilbert O Ouma, Mohamed Y Said, Gordon O Wayumba, and Joseph Muhwang. Spatial and temporal trends of rainfall and temperature in the Amboseli ecosystem of Kenya. Technical University of Kenya Institutional Repository, 5 (5):28–42, 2018. URL http://repository.tukenya.ac.ke/handle/123456789/1828.

[47] Susan C Alberts and Jeanne Altmann. Immigration and hybridization patterns of yellow and anubis baboons in and around Amboseli, Kenya. American Journal of Primatology, 53(4):139–154, 2001. ISSN 0275-2565. URL https://onlinelibrary.wiley.com/doi/abs/10.1002/ajp.1.

[48] Tauras P. Vilgalys, Arielle S. Fogel, Jordan A. Anderson, Raphael S. Mututua, J. Kinyua Warutere, I. Long’ida Siodi, Sang Yoon Kim, Tawni N. Voyles, Jacqueline A. Robinson, Jeffrey D. Wall, Elizabeth A. Archie, Susan C. Alberts, and Jenny Tung. Selection against admixture and gene regulatory divergence in a long-term primate field study. Science, 377(6606):635–641, 2022. doi: 10.1126/science.abm4917. URL https://www.science.org/doi/abs/10.1126/science.abm4917.

[49] Arielle S. Fogel, Peter O. Oduor, Albert W. Nyongesa, Charles N. Kimwele, Susan C. Alberts, Elizabeth A. Archie, and Jenny Tung. Ecology and age, but not genetic ancestry, predict fetal loss in a wild baboon hybrid zone. American Journal of Biological Anthropology, 180(4):618–632, 2023. ISSN 2692-7691. doi: 10.1002/ajpa.24686. URL https://onlinelibrary.wiley.com/doi/abs/10.1002/ajpa.24686.

[50] Stuart A Altmann. Foraging for survival: yearling baboons in Africa. University of Chicago Press, 1998. ISBN 0226015955.

[51] Linda Marie Fedigan, Sarah D. Carnegie, and Katharine M. Jack. Predictors of reproductive success in female white-faced capuchins (cebus capucinus). American Journal of Physical Anthropology, 137(1):82–90, 2008. ISSN 0002-9483. doi: 10.1002/ajpa.20848. URL https://onlinelibrary.wiley.com/doi/abs/10.1002/ajpa.20848.

[52] Emily J. Levy, Laurence R. Gesquiere, Emily McLean, Mathias Franz, J. Kinyua Warutere, Serah N. Sayialel, Raphael S. Mututua, Tim L. Wango, Vivian K. Oudu, Jeanne Altmann, Elizabeth A. Archie, and Susan C. Alberts. Higher dominance rank is associated with lower glucocorticoids in wild female baboons: A rank metric comparison. Hormones and Behavior, 125:104826, 2020. ISSN 0018-506X. doi: 10.1016/j.yhbeh.2020.104826. URL http://www.sciencedirect.com/science/article/pii/S0018506X20301525.

[53] Jacob B. Gordon, David Jansen, Niki H. Learn, Jeanne Altmann, Jenny Tung, Elizabeth A. Archie, and Susan C. Alberts. Ordinal dominance rank assignments: Protocol for the Amboseli Baboon Research Project, 2023. URL https://amboselibaboons.nd.edu/assets/509243/gordon_etal_elo_matrix_rank_comparisons_22mar2023_for_abrp_website_google_docs.pdf.

[54] Laurence R. Gesquiere, Jeanne Altmann, Elizabeth A. Archie, and Susan C. Alberts. Interbirth intervals in wild baboons: Environmental predictors and hormonal correlates. American Journal of Physical Anthropology, 166(1):107–126, 2018. ISSN 0002-9483. doi: 10.1002/ajpa.23407. URL https://onlinelibrary.wiley.com/doi/abs/10.1002/ajpa.23407.

[55] Jeanne Altmann, Laurence Gesquiere, Jordi Galbany, Patrick O. Onyango, and Susan C. Alberts. Life history context of reproductive aging in a wild primate model. Annals of the New York Academy of Sciences, 1204(1):127–138, 2010. ISSN 0077-8923. doi: 10. 1111/j.1749-6632.2010.05531.x. URL https://nyaspubs.onlinelibrary.wiley.com/doi/abs/10.1111/j.1749-6632.2010.05531.x.

[56] Jeanne Altmann and Susan C. Alberts. Variability in reproductive success viewed from a life-history perspective in baboons. American Journal of Human Biology, 15(3):401– 409, 2003. ISSN 1042-0533. doi: 10.1002/ajhb.10157. URL https://onlinelibrary.wiley.com/doi/abs/10.1002/ajhb.10157.

[57] Emily M. McLean, Elizabeth A. Archie, and Susan C. Alberts. Lifetime fitness in wild female baboons: Trade-offs and individual heterogeneity in quality. The American Naturalist, 194(6):745–759, 2019. doi: 10.1086/705810. URL https://www.journals.uchicago.edu/doi/abs/10.1086/705810.

[58] Joshua D Angrist and Jorn-Steffen Pischke. Mostly Harmless Econometrics. Princeton University Press, Princeton, NJ, 2008. ISBN 1400829828.

[59] Jeffery M. Wooldridge. Introductory Econometrics: A Modern Approach. Cengage Learning, Boston, MA, 6th edition, 2016.

[60] Panchanan Das. Time series: data generating process, pages 247–259. Springer, Singapore, 2019. ISBN 9813290188.

[61] Olive Jean Dunn. Multiple comparisons among means. Journal of the American Statistical Association, 56(293):52–64, 1961. ISSN 0162-1459. doi: 10.1080/01621459.1961. 10482090. URL https://www.tandfonline.com/doi/abs/10.1080/01621459.1961.10482090.

[62] Tyler J VanderWeele and Maya B Mathur. Some desirable properties of the Bonferroni correction: Is the Bonferroni correction really so bad? American Journal of Epidemiology, 188(3):617–618, 2018. ISSN 0002-9262. doi: 10.1093/aje/kwy250. URL 10.1093/aje/kwy250.

[63] Yoav Benjamini and Yosef Hochberg. Controlling the false discovery rate: A practical and powerful approach to multiple testing. Journal of the Royal Statistical Society: Series B, 57(1):289–300, 1995. ISSN 0035-9246. doi: 10.1111/j.2517-6161.1995.tb02031.x. URL https://rss.onlinelibrary.wiley.com/doi/abs/10.1111/j.2517-6161.1995.tb02031.x.

[64] Michael L. Anderson. Multiple inference and gender differences in the effects of early intervention: A reevaluation of the abecedarian, perry preschool, and early training projects. Journal of the American Statistical Association, 103(484):1481–1495, 2008. ISSN 01621459. URL http://www.jstor.org/stable/27640197.

[65] Marco Festa-Bianchet, Jon T. Jorgenson, and Denis Ŕeale. Early development, adult mass, and reproductive success in bighorn sheep. Behavioral Ecology, 11(6):633–639, 2000. ISSN 1045-2249. doi: 10.1093/beheco/11.6.633. URL 10.1093/beheco/11.6.633.

[66] Morgane Gicquel, Marion L. East, Heribert Hofer, and Sarah Benhaiem. Early-life adversity predicts performance and fitness in a wild social carnivore. Journal of Animal Ecology, 91(10):2074–2086, 2022. ISSN 0021-8790. doi: 10.1111/1365-2656.13785. URL https://besjournals.onlinelibrary.wiley.com/doi/abs/10.1111/1365-2656.13785.

[67] Robin E Morrison, Winnie Eckardt, Fernando Colchero, Veronica Vecellio, and Tara S Stoinski. Social groups buffer maternal loss in mountain gorillas. Elife, 10:e62939, 2021. ISSN 2050-084X. URL https://elifesciences.org/articles/62939.

[68] Phyllis C Lee, Cynthia J Moss, Norah Njiraini, Joyce H Poole, Katito Sayialel, and Vicki L Fishlock. Cohort consequences of drought and family disruption for male and female African elephants. Behavioral Ecology, 33(2):408–418, 2021. ISSN 1465-7279. doi: 10.1093/beheco/arab148. URL 10.1093/beheco/arab148.

[69] Harry H Marshall, Emma IK Vitikainen, Francis Mwanguhya, Robert Businge, Solomon Kyabulima, Michelle C Hares, Emma Inzani, Gladys Kalema-Zikusoka, Kenneth Mwesige, and Hazel J Nichols. Lifetime fitness consequences of early-life ecological hardship in a wild mammal population. Ecology and Evolution, 7(6):1712–1724, 2017. ISSN 2045-7758. URL https://onlinelibrary.wiley.com/doi/full/10.1002/ece3.2747.

[70] Theresa M Lee and Irving Zucker. Vole infant development is influenced perinatally by maternal photoperiodic history. *American Journal of Physiology-Regulatory*, Integrative and Comparative Physiology, 255(5):R831–R838, 1988. ISSN 0363-6119. URL https://journals.physiology.org/doi/abs/10.1152/ajpregu.1988.255.5.R831.

[71] Ben Dantzer, Amy E. M. Newman, Rudy Boonstra, Rupert Palme, Stan Boutin, Murray M. Humphries, and Andrew G. McAdam. Density triggers maternal hormones that increase adaptive offspring growth in a wild mammal. Science, 340(6137):1215–1217, 2013. doi: 10.1126/science.1235765. URL https://www.science.org/doi/abs/10.1126/science.1235765.

[72] Fernando A. Campos, Francisco Villavicencio, Elizabeth A. Archie, Fernando Colchero, and Susan C. Alberts. Social bonds, social status and survival in wild baboons: a tale of two sexes. Philosophical Transactions of the Royal Society B: Biological Sciences, 375(1811):20190621, 2020. doi: 10.1098/rstb.2019.0621. URL https://royalsocietypublishing.org/doi/abs/10.1098/rstb.2019.0621.

[73] Lauren Petrullo, David Delaney, Stan Boutin, Jeffrey E. Lane, Andrew G. McAdam, and Ben Dantzer. A future food boom rescues the negative effects of cumulative early-life adversity on adult lifespan in a small mammal. bioRxiv, page 2023.08.16.553597, 2023. doi: 10.1101/2023.08.16.553597. URL https://www.biorxiv.org/content/biorxiv/early/2023/11/28/2023.08.16.553597.full.pdf.

[74] Wolfgang P. J. Dittus. Group fission among wild toque macaques as a consequence of female resource competition and environmental stress. Animal Behaviour, 36(6): 1626–1645, 1988. ISSN 0003-3472. doi: 10.1016/S0003-3472(88)80104-0. URL https://www.sciencedirect.com/science/article/pii/S0003347288801040.

[75] Brian A. Lerch, Karen C. Abbott, Elizabeth A. Archie, and Susan C. Alberts. Better baboon break-ups: collective decision theory of complex social network fissions. Proceedings of the Royal Society B: Biological Sciences, 288(1964):20212060, 2021. doi: 10.1098/rspb.2021.2060. URL https://royalsocietypublishing.org/doi/abs/10.1098/rspb.2021.2060.

[76] Mathieu Douhard, Floriane Plard, Jean-Michel Gaillard, Gilles Capron, Daniel Delorme, Fraņcois Klein, Patrick Duncan, Leif Egil Loe, and Christophe Bonenfant. Fitness consequences of environmental conditions at different life stages in a long-lived vertebrate. Proceedings of the Royal Society B: Biological Sciences, 281(1785):20140276, 2014. doi: doi:10.1098/rspb.2014.0276. URL https://royalsocietypublishing.org/doi/abs/10.1098/rspb.2014.0276.

[77] Chelsea J. Weibel, Jenny Tung, Susan C. Alberts, and Elizabeth A. Archie. Accelerated reproduction is not an adaptive response to early-life adversity in wild baboons. Proceedings of the National Academy of Sciences, 117(40):24909–24919, 2020. doi: 10.1073/pnas.2004018117. URL https://www.pnas.org/content/pnas/117/40/24909.full.pdf.

[78] Margaret A. Stanton, Elizabeth V. Lonsdorf, Carson M. Murray, and Anne E. Pusey. Consequences of maternal loss before and after weaning in male and female wild chimpanzees. Behavioral Ecology and Sociobiology, 74(2):22, 2020. ISSN 1432-0762. doi: 10.1007/s00265-020-2804-7. URL 10.1007/s00265-020-2804-7.

[79] Mirkka Lahdenpera, Khyne U. Mar, and Virpi Lummaa. Short-term and delayed effects of mother death on calf mortality in Asian elephants. Behavioral Ecology, 27(1):166– 174, 2015. ISSN 1045-2249. doi: 10.1093/beheco/arv136. URL 10.1093/beheco/arv136.

[80] Eli D. Strauss, Daizaburo Shizuka, and Kay E. Holekamp. Juvenile rank acquisition is associated with fitness independent of adult rank. Proceedings of the Royal Society B: Biological Sciences, 287(1922):20192969, 2020. doi: 10.1098/rspb.2019.2969. URL https://royalsocietypublishing.org/doi/abs/10.1098/rspb.2019.2969.

